# SpaFun: Discovering Domain-specific Spatial Expression Patterns and New Disease-Relevant Genes using Functional Principal Component Analysis

**DOI:** 10.1101/2025.02.17.638766

**Authors:** Xi Jiang, Yanghong Guo, Lei Guo, Lin Zhong, Jiayi Wang, Guanghua Xiao, Qiwei Li, Lin Xu

**Affiliations:** Quantitative Biomedical Research Center, Peter O’Donnell Jr. School of Public Health, The University of Texas Southwestern Medical Center, Dallas, Texas, U.S.A; Department of Statistics and Data Science, Southern Methodist University, Dallas, Texas, U.S.A.; Department of Mathematical Sciences, The University of Texas at Dallas, Richardson, Texas, USA

**Keywords:** domain-representative gene (DRG), functional principal component analysis (fPCA), spatial expression pattern, spatially resolved transcriptomics (SRT), spatially variable gene (SVG)

## Abstract

SpaFun is a novel, non-model-based method developed to address limitations in existing spatially variable gene (SVG) detection techniques, particularly for large-scale spatially resolved transcriptomics (SRT) datasets. These limitations include computational inefficiency, limited statistical power with increasing data size, and the inability to capture spatial heterogeneity and co-expression patterns among genes. Built on functional principal component analysis (fPCA), SpaFun identifies domain-representative genes (DRGs) with significantly better computational efficiency and greater statistical power while accounting for spatial heterogeneity and co-expression patterns among genes. We applied SpaFun to three SRT datasets and demonstrated that SpaFun outperformed state-of-the-art algorithms for identifying representative genes for tumor regions (e.g., DESeq, edgeR, and limma), as well as recently developed novel algorithms designed for spatial omics to identify the representative genes (e.g., SPARK and CSIDE). This highlights SpaFun’s ability to accurately identify genes most representative of each spatial domain (e.g., tumor, immune, or stroma regions). By uncovering novel disease-relevant genes overlooked by existing algorithms, SpaFun could provide insights into new molecular mechanisms and propose innovative therapeutic strategies to improve patient outcomes.

**Key Points:**

- SpaFun is the first method dedicated to identifying DRGs, capturing spatially representative expression patterns within annotated tissue regions, setting it apart from all traditional SVG detection methods.
- It leverages fPCA to model gene expression as a function of spatial location, avoiding reliance on predefined spatial distribution assumptions.
- The non-model-based framework ensures compatibility with different SRT platforms and experimental designs, making it a scalable and widely applicable tool for SRT research.

## INTRODUCTION

Spatially resolved transcriptomics (SRT), a cutting-edge technique that has emerged in recent years, provides gene expression measurements while incorporating spatial information. This enables the study of various biological phenomena, such as cell-cell communication, cell heterogeneity, and regulation of transcriptional networks. There are two types of SRT technologies: imaging-based and sequencing-based spatial omics. Imaging-based methods, such as STARmap [1], MERFISH [2], and seqFISH [3], utilize combinatorial labeling and sequential imaging to achieve subcellular visualization and quantification of transcripts. However, they are limited by low throughput on the number of genes studied in each experiment, high cost, and complexity of experimental designs [4–7]. In contrast, sequencing-based techniques such as Spatial transcriptomics (ST), Visium, and Slide-seq, provide higher throughput for analyzing large numbers of genes and ease of use, albeit at the cost of limited capability to achieve single-cell resolution [8–10].

Based on the advantages and limitations of these techniques, researchers have developed a variety of methods for analyzing SRT data. A crucial type of analysis is to detect spatially variable genes (SVGs), which hold promise for identifying important biomarkers in pathological processes and embryonic development. Unlike highly variable gene (HVG) detection algorithms [11–13], which focus on identifying genes with higher variability among spots/cells, the SVG detection method aims to discover genes with unique spatial expression patterns but not necessarily highly variable. While various methods have been developed to identify SVGs, most of them focus on detecting genes with spatial patterns across the entire tissue encompassing different histological regions. For example, statistical-modeling-based methods such as SpatialDE [14], SPARK [15], GPcounts [16], and BoostGP [17] utilize Gaussian processes (GPs) to model variations in spatial patterns in transcriptomic expression. As modeling assumptions may not always hold, alternative approaches without covariance kernel setup requirements could facilitate improved spatially variable gene identification. Grid-based strategies often partition the 2-D space into grids and define neighbors for each cell or spot. For instance, singleCellHaystack [18] computes a local reference distribution around a fixed cell set for a target gene, enabling the derivation of KL-divergence and p-values through comparison against the null hypothesis of uniform distribution across cells. Similarly, MERINGUE [19] identifies SVGs using Moran’s I. An alternative approach, BinSpect [20], constructs spatial grids and dichotomizes gene expression into high versus low to assess the correlation of gene expression levels among neighboring cells. Machine-learning methods like SpaGCN [21] first identify spatial domains (i.e., regions with similar spatial expression patterns) based on gene expression and histological characteristics, followed by the detection of highly expressed genes within specific domains using the Wilcoxon rank-sum test. These methods are often computationally intensive when the dimension of the input gene expression data is high.

As SRT technology has advanced, it has become increasingly common to generate large-scale SRT datasets, which offer unprecedented insights into the spatial organization of gene expression at a high resolution. Existing SVG detection methods may suffer from limited applicability for large-scale datasets due to their high computational complexity. In addition, most of the existing methods focus on identifying gene expression patterns on the entire spatial landscape, which can overlook spatial heterogeneity and result in lower statistical power as the number of measured sample points increases. Moreover, most current methods identify SVGs independently for each gene, neglecting the potential spatial contexts, co-expression patterns, and interaction among genes. These limitations of existing SVG detection methods motivate us to introduce functional principal component analysis (fPCA) into this area, aiming at developing a domain-representative gene (DRG) detection method. fPCA is a statistical method used to analyze and reduce the dimensionality of functional data. It identifies the main modes of variation within the data, helping to capture the most important patterns or trends across the entire functional dataset. Similar to SVGs, DRGs exhibit spatial patterns within regions; however, DRGs also learn unique, representative patterns specific to each spatial domain, providing a more detailed understanding of spatially organized gene expression.

In this study, we propose SpaFun, a non-model-based approach built upon fPCA to identify DRGs with spatial patterns within a specific annotated region (i.e., an annotated spatial domain), particularly the tumor region. With the help of fPCA, this method avoids specific assumptions about spatial distributions of gene expression, unlike Gaussian process-based or grid-based approaches. It considers gene expression as a function of its spatial location and aims to detect representative domain-specific spatial expression patterns. Corresponding DRGs for each spatial expression pattern are then identified. We applied SpaFun to three SRT datasets, two generated from sequencing-based (e.g., 10X Visium) and one from imaging-based (e.g., STARmap) platforms. The results demonstrate that SpaFun can identify biologically meaningful DRGs based on distinct spatial omics technologies. By focusing on detecting spatial patterns within defined regions and accounting for interaction among genes, SpaFun enhances the understanding of localized gene expression dynamics, contributing to a deeper comprehension of complex biological processes.

## METHODS

In this section, we first define the histology profile, which includes domain information, as well as geospatial and molecular profiles derived from SRT data. We then discuss the two stages of SpaFun: (1) identifying eigen-spatial expression patterns using functional principal component analysis (fPCA) and (2) detecting domain-representative genes (DRGs) for each identified spatial expression pattern.

### Data preparation

#### Histology profile *z*

To deal with the heterogeneity in cellular spatial organization, we partition the whole domain into *K* mutually exclusive histology-based spatial domains as an initial step. We use ***Z*** = (*z*_1_, ⋯, *z*_*N*_)^T^ to denote the vector specifying the identity of the *K* domains. The spatial domains ***Z*** is decided by prior biological or clinical knowledge, or estimated by existing methods based on molecular profile or/and histology image. In our analysis of 10x Visium ST data, we use the manual annotation from the pathologist. The layer structure from the original study is applied to define domains for the mouse visual cortex STARmap data.

For our method, we focus on a specific spatial domain and denote the target domain as *k*. Thus, we extract the SRT data for the following analysis by selecting the spots within this specific domain.

#### Geospatial profile *T*

For NGS-based techniques, spots are the round area of barcoded mRNA capture probes where gene expression levels are measured. The spatial distribution of spots is arrayed on a square or triangular lattice. We denote the SRT geospatial profile by an *N* × 2 matrix ***T***, where each row ***t***_***i***_ = (*t*_*i*1_, *t*_*i*2_) gives the *x* and *y* coordinates of the spot *i* on a two-dimensional Cartesian plane. ST and 10x Visium spots are arranged on square and triangular lattice grids, respectively. For imaging-based SRT platforms, they are able to capture the gene expression at single-cell resolution. Similarly, we use matrix ***T*** to denote the locations of cells, where each row ***t***_***i***_ = (*t*_*i*1_, *t*_*i*2_) gives the *x* and *y* coordinates of the cell *i*. For the geospatial profile in the target domain *k*, we extract the rows of matrix ***T*** with spot/cell *i* belongs to this domain (i.e., *z*_*i*_ = *k*) and denote the matrix as ***T***_*k*_.

#### Molecular profile *C*

In general, the spot-level molecular profile of NGS-based SRT data can be represented by an *N* × *P* count table ***C***, where each entry *c*_*ij*_ ∈ ℕ, *i* = 1, ⋯, *N*, *j* = 1, ⋯, *P* is the read count for gene *j* measured at spot *i*. For the target domain, similarly to the geospatial profiles, we extract the rows of the count table with corresponding spots/cells *i* in this domain. We denote the extracted SRT data for a target domain *k* as ***C***_*k*_, an *N*_*k*_ × *P* count matrix, where *N*_*k*_ is the number of spots/cells in a domain *k* (i.e., 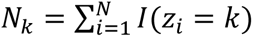). Count matrix ***C***_*k*_ is then normalized and transformed to relative expression levels by following the normalization steps of SpatialDE. First, we use a variance-stabilizing transformation for negative-binomial-distributed data known as Anscombe’s transformation to obtain a relatively normally distributed expression level vector for each gene. Second, to account for those nuisance effects across spots, including sequencing depth, amplification, dilution efficiency, and reverse transcription efficiency, we regress log total count values out from the Anscombe-transformed expression values before fitting the spatial models. We denote the normalized gene expression matrix as ***C̃***_*k*_. For a particular gene *j*, the expression level vector is denoted as ***c̃***_*k*,*j*_.

### Stage I: Identify eigen-spatial expression patterns by fPCA

It is very common to use eigen-images to identify dominant spatial patterns of variation based on data observed at *N* locations with *P* samples. A typical approach to estimate eigen-images is principal component analysis (PCA). However, when the number of samples *P* is large compared to the number of observed locations *N* (in our case, *P* is approximately 10,000 v.s. *N* is approximately 1,000), PCA may lead to noisy eigen-images which do not provide meaningful insights. To alleviate this issue, people usually put a smooth assumption on eigen-images to seek for continuous spatially localized patterns. Following this assumption, we propose to identify eigen-spatial expression patterns by functional principal component analysis (fPCA) for every (compact) domain *D*_*k*_ ∈ ℝ^2^, *k* = 1, … *K*.

In the view of functional data, we take {*X*_*k*,*j*_(***t***), ***t*** ∈ *D*_*k*_, *j* = 1, …, *p*} as independent copies of zero-mean ℒ^2^ -continuous stochastic process *X*_*k*_ defined on the domain *D*_*k*_. And we have ***c̃***_*k*,*j*,*i*_ = *X*_*k*,*j*_(***t***_*k*,*i*_) + *∈*_*k*,*j*,*i*_ for *j* = 1, ⋯, *P* and *i* = 1, ⋯, *N*, where ***c̃***_*k*,*j*,*i*_ is the *i*-th entry of vector ***c̃***_*k*,*j*_, ***t***_*k*,*i*_ is the *i*-th row of matrix ***T***_*k*_, and {*∈*_*k*,*j*,*i*_, *j* = 1, ⋯, *P*, *i* = 1, ⋯, *N*} are independent errors with zero mean and finite variance. Take Γ_*k*_ as the covariance function of *X*_*k*_, which is defined as Γ_*k*_(***t***, ***t***′) = *Cov*(*X*_*k*_(***t***), *X*_*k*_(***t***′)). By mercer’s theorem, we can represent the covariance function in terms of the spectral decomposition via the ℒ^2^ inner product: Γ_*k*_(***t***, ***t***′) = ∑_*l*≥1_ ζ_*k*,*l*_*ϕ*_*k*,*l*_ (***t***)*ϕ*_*k*,*l*_(***t***′), where ζ_*k*,1_ ≥ ζ_*k*,2_ ≥ ⋯ ≥ 0 are eigenvalues and {*ϕ*_*k*,*l*_, *l* = 1, …, ∞ } are eigenfunctions which have the following properties: 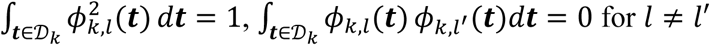. Then, to obtain eigenfunctions (eigen-images) that show the dominant spatial patterns, we only need to estimate the covariance function Γ_*k*_.

In practice, to promote dimension reduction and enhance interoperability, we are interested in obtaining a low-rank estimation of Γ_*k*_, i.e., there are finite numbers of eigenfunctions. As such, we adopt the idea of reproducing kernel Hilbert space (RKHS) modeling of the covariance function proposed in [22] and [23] to obtain a smooth and low-rank estimation of Γ_*k*_.

We assume the sample processes of *X*_*k*_ lie in an RKHS ℋ_*k*_ of functions defined on *D*_*k*_ with a continuous and square integrable reproducing kernel *K*. Let ⟨⋅,⋅⟩_ℋ*k*_ and ||⋅ ||_ℋ*k*_ denote the inner product and norm of ℋ_*k*_ respectively. Then we have Γ_*k*_ ∈ ℋ_*k*_ ⊗ ℋ_*k*_, where ℋ_*k*_ ⊗ ℋ_*k*_ is again an RKHS with reproducing kernel *K̃*((***s***, ***t***), (***s***′, ***t***′)) = *K*(***s***, ***s***′)*K*(***t***, ***t***′), under the technical condition 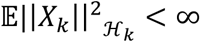. Define ℳ = {Γ ∈ ℋ_*k*_ ⊗ ℋ_*k*_, Γ *is positive definite*}. We estimate the Γ_*k*_ via the following optimization:

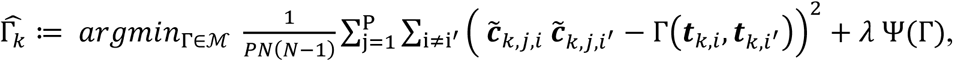

Where *λ* > 0 is a tuning parameter and Ψ(⋅) is the trace-norm regularization function that promotes the low-rankness. See details of RKHS framework of functional data and detailed definition of Ψ(⋅) in Section 2 and definition 5 of [23], respectively.

After obtaining 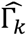 with an appropriately chosen kernel *K*, we adopt the procedure described in Section 2.5 in [22] to get the estimation of eigenfunctions 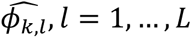. Eigenfunctions with variance explained greater than 10% are selected. The selected eigenfunctions 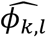 and their reflection over the x-axis –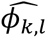 are then defined as representative spatial expression patterns. We applied Gaussian kernel in this study, where the bandwidth was chosen by median heuristic [24].

### Stage II: Detect domain representative genes (DRGs)

After we obtain the top *l* representative spatial expression patterns for domain *k*, we identify the DRGs corresponding to each spatial pattern by calculating the similarity between eigenfunctions and expression levels of individual genes. Each eigenfunction represents a PC. To measure similarity, we perform a Pearson correlation test for each pair of PC and gene. This test provides a correlation coefficient and a corresponding p-value that quantifies the similarity between the given pair. To account for multiple comparisons, adjusted p-values are calculated using the Benjamini-Hochberg procedure, which controls the false discovery rate (FDR) across all genes for each PC, providing a more accurate significance level. In the real data analysis presented in this paper, we define DRGs as genes with an absolute correlation coefficient > 0.3 and an adjusted p-value < 0.01.

## RESULTS

### Overview of SpaFun

SpaFun is a non-model-based approach that leverages functional principal component analysis (fPCA) to identify DRGs with spatial expression patterns in specific regions, such as tumor areas (as shown in Figure 1). Unlike existing methods, SpaFun efficiently handles large-scale SRT datasets, accounting for spatial heterogeneity, and captures co-expression among genes. We applied SpaFun, along with other comparable methods for detecting HVGs and SVGs, to three real datasets: human prostate cancer, human ovarian cancer, and mouse visual cortex STARmap [1], derived from various spatial omics platforms. Downstream analysis, such as Gene Ontology (GO) enrichment analysis, further demonstrated that SpaFun effectively detected DRGs that align closely with known biological knowledge. By uncovering biologically meaningful DRGs, SpaFun advances our understanding of localized gene expression, providing a detailed characterization of the SRT landscape in complex tissues.

**Figure 1.**
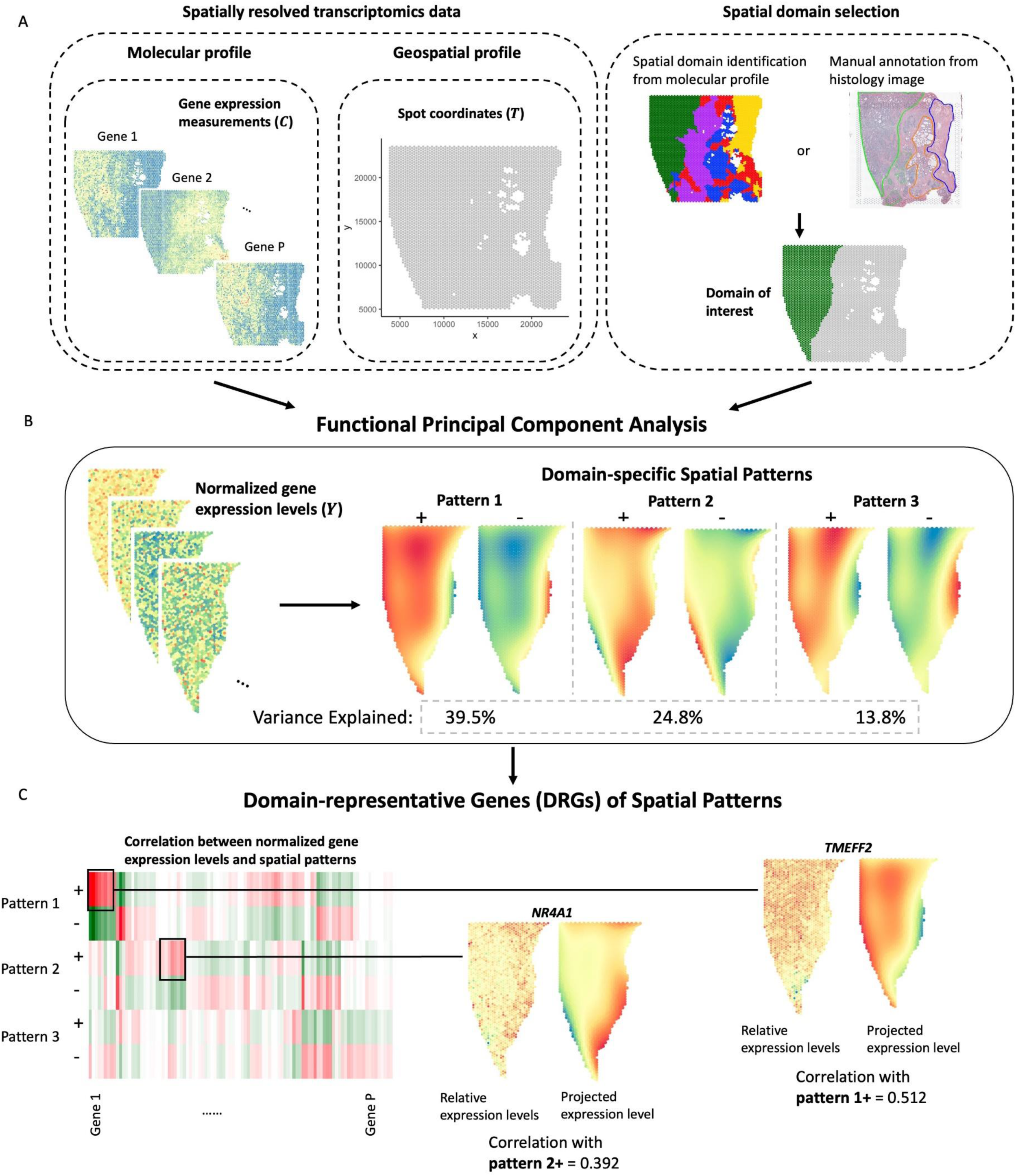
Flowchart of SpaFun. SpaFun is a non-model-based approach that utilizes functional principal component analysis (fPCA) to identify domain representative genes (DRGs) with spatial expression patterns in specific regions. (**A**) Gene expression data and spatial information serve as inputs for SpaFun, with a pre-selected spatial domain for analysis. (**B**) fPCA is performed on the selected domain, and the top domain-specific spatial patterns, determined by the leading principal components, are chosen as representative patterns for that domain. (**C**) DRGs are identified based on each domain-specific spatial pattern.

### Application to human prostate cancer dataset

We applied SpaFun to analyze an SRT dataset from a human prostate cancer study, comprising 4,371 spots and 17,651 genes. Gene expression was measured using the 10x Visium platform on a tissue section from an invasive carcinoma of the prostate. This dataset includes manual annotations provided by pathologists, as displayed in Figure 2A. Three annotated tissue regions are identified: tumor, stroma, and stroma & partially atrophic changes. The tumor domain, highlighted in Figure 2B, is of primary interest (results on the stroma region is provided in Figure S1 in the supplementary information).

**Figure 2.**
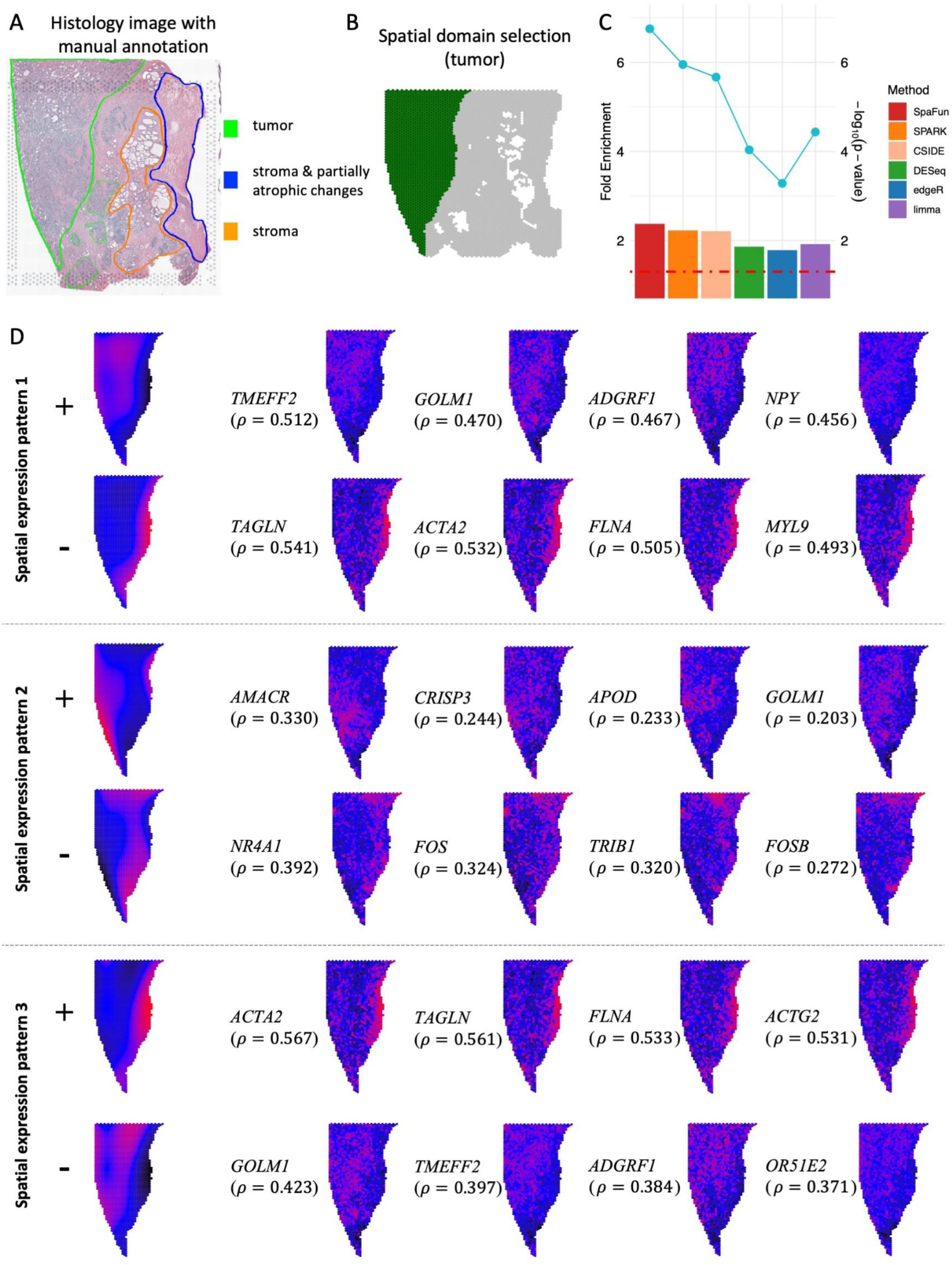
Results of the human prostate cancer dataset. (**A**) Histology image with pathologist-annotated spatial domains. (**B**) Tumor domain selected for analysis. (**C**) Gene set enrichment analysis results for six methods: SpaFun, SPARK, CSIDE, DESeq, edgeR, and limma. Bar plots represent the fold enrichment of each method, with the dotted blue line indicating the −log_10_(p-value) for each method and the dashed red horizontal line marking the threshold where p-value = 0.05. (**D**) The top three principal components (PCs) in the tumor domain, each shown in two opposite directions, along with the corresponding genes exhibiting similar expression patterns to each PC.

Figure 2D depicts the top three principal components (PCs) in the tumor domain, where each PC represents a distinct spatial expression pattern. Taking the first PC as an example, it is important to note that, as with all PCA methods, each PC can have two opposite directions. While the direction affects the color coding, it does not alter the spatial expression pattern. For simplicity, we focus on the positive direction as an example. In this visualization, black indicates low expression levels, red indicates high expression levels, and blue represents intermediate levels. The first PC shows higher expression in the middle region and lower expression along the right edge, with the opposite pattern in the negative direction. For both directions, genes with expression patterns similar to the PC can be identified. For example, TMEFF2 and GOLM1 exhibit higher expression in the center and lower expression on the right edge, while TAGLN and ACTA2 show the opposite trend, with lower expression in the center and higher expression on the right edge. As illustrated in Figure 2D, each PC highlights a unique spatial expression pattern.

The results of gene set enrichment analysis are presented in Figure 2C. The dotted blue line represents the −log_10_(p-value) for each method, while the dashed horizontal red line indicates the threshold where p-value = 0.05. The accompanying barplot illustrates the fold enrichment across all comparable methods. Both the p-value and fold enrichment analyses demonstrate the effectiveness of SpaFun in identifying biologically meaningful genes within the prostate tumor domain. DESeq, edgeR, and limma are widely recognized methods for identifying differentially expressed genes to represent the omics features of each spatial domain. Meanwhile, SPARK and CSIDE are recent algorithms specifically designed for spatial omics datasets. In this analysis, SpaFun outperformed all these methods, demonstrating its superior ability to accurately identify the genes most representative of each spatial domain.

We conducted GO enrichment analysis on the selected DRGs for each PC to investigate their associated biological functions. A summary of the GO enrichment analysis, highlighting the enriched expression patterns for each PC, is presented in Figure S2 in the supplementary information. For each PC, genes with p-values < 0.01 from the correlation test with the PC were selected as DRGs. Our analysis identified 1,247, 86, 1,153, 529, and 48 enriched GO terms associated with the first five PCs, respectively, using an adjusted p-value threshold of 0.05. The top 15 GO terms with the smallest p-values for the first five PCs are displayed in Figure S2.

### Application to human ovarian cancer dataset

The second SRT dataset we analyzed in this study is derived from a section of human ovarian tumor tissue. This dataset includes 3,455 spots and 17,651 genes. Gene expression was measured on a section of serous papillary carcinoma from human ovarian tissue using the 10x Visium platform, with the H&E-stained image shown in Figure 3A. Tumor and stroma domains were annotated by pathologists. Since the tumor domain is of primary interest (Figure 3B), the results for this domain are presented in Figure 3, while the results for the stroma domain are provided in Figure S3 in the supplementary information.

**Figure 3.**
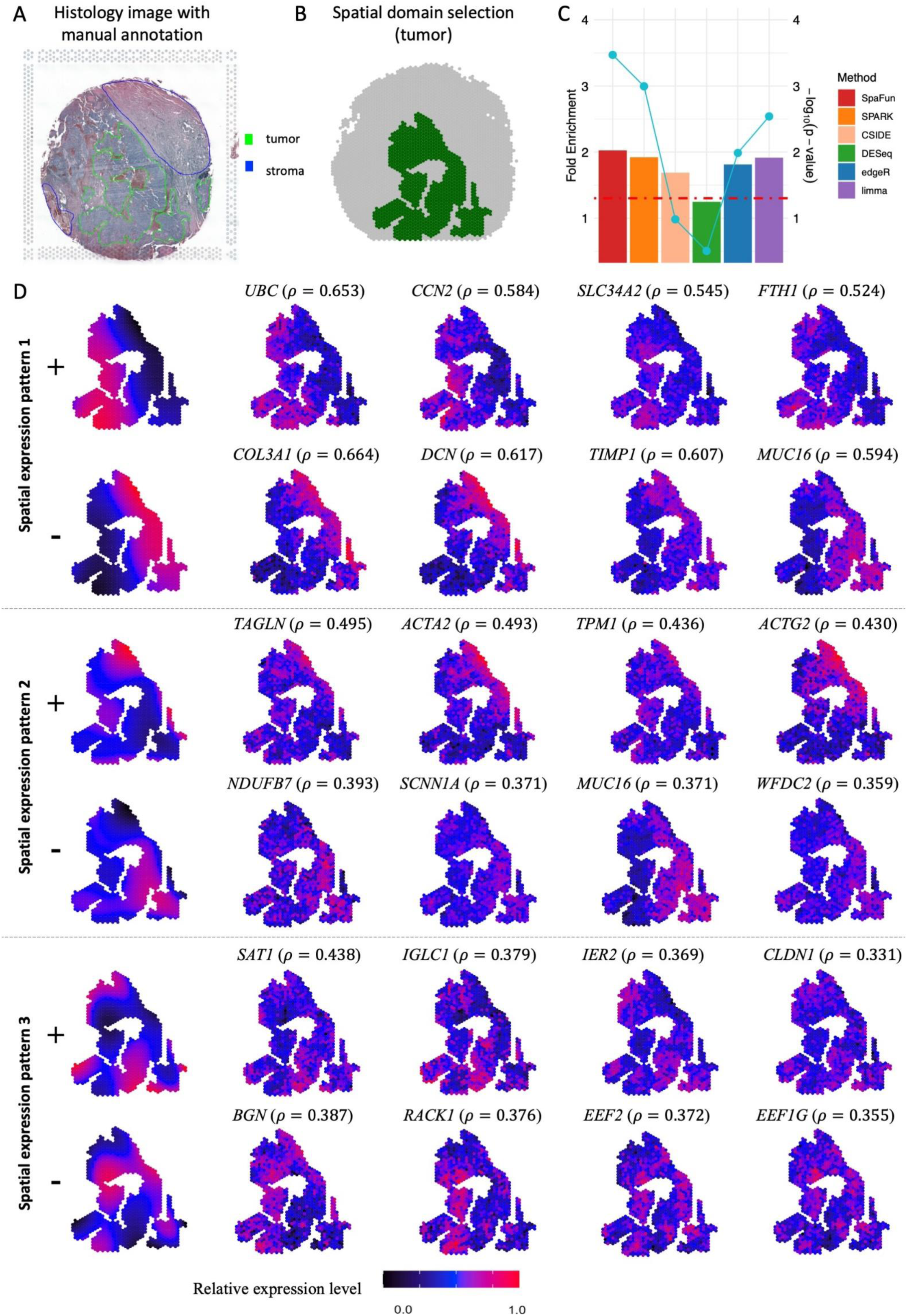
Result of the human ovarian cancer dataset. (**A**) Histology image with pathologist-annotated spatial domains. (**B**) Tumor domain selected for analysis. (**C**) Gene set enrichment analysis results for six methods: SpaFun, SPARK, CSIDE, DESeq, edgeR, and limma. Bar plots represent the fold enrichment of each method, with the dotted blue line indicating the −log_10_(p-value) for each method and the dashed red horizontal line marking the threshold where p-value = 0.05. (**D**) The top three principal components (PCs) in the tumor domain, each shown in two opposite directions, along with the corresponding genes exhibiting similar expression patterns to each PC.

Each PC represents a unique expression pattern, and Figure 3D illustrates the top three PCs along with genes that share similar expression patterns. Taking PC1 as an example, the positive direction of this PC shows high expression levels in the bottom-left corner, which gradually decreases along the bottom-up diagonal direction. Conversely, the negative direction exhibits the opposite pattern. By calculating the correlation between each gene and the PC, we identified genes with the highest correlation coefficients, as shown in the figure. For instance, UBC and CCN2 display expression patterns with higher levels in the bottom-left region and lower levels in the top-right, while COL3A1 and DCN exhibit the reverse pattern and are highly correlated with the negative direction of PC1.

The results of six comparable methods are shown in Figure 3C. The dotted blue line represents the −log_10_(p-value) for each method, while the dashed horizontal red line indicates the threshold where p-value =0.05. Four methods—SpaFun, SPARK, edgeR, and limma—achieved a p-value < 0.05, whereas the other two methods, CSIDE and DESeq, had p-values >0.05. The accompanying bar plot illustrates the fold enrichment across all methods. Among the six, SpaFun demonstrated the lowest p-value and the highest fold enrichment compared with the other state-of-the-art methods, highlighting its effectiveness in detecting biologically meaningful genes in the prostate tumor domain.

We conducted GO enrichment analysis on the selected DRGs for each PC to investigate their associated biological functions. Figure S4 in the supplementary information provides a summary of the GO enrichment analysis and highlights the enriched expression patterns for each PC. For each PC, genes with a p-value < 0.01 from the correlation test with the PC were selected as DRGs. Our analysis identified 1,076, 1,280, and 476 enriched GO terms associated with the first three PCs, respectively, using an adjusted p-value threshold of 0.05. No enriched terms were found for PCs beyond the first three. The top 15 GO terms with the smallest p-values for the first three PCs are presented in Figure S4.

### Application to mouse visual cortex STARmap data

To demonstrate that SpaFun is also applicable to single-cell-level SRT data, we applied it to a STARmap dataset. This dataset was generated from the mouse visual cortex, including the hippocampus, corpus callosum, and neocortical layers. In total, 1020 genes were measured across 1207 cells representing 15 cell types. The detailed structure of the layers is shown in Figure 4A. Layer L4 was selected for the comparison experiment, while the results for L2/3 are provided in Figure S5 of the supplementary information.

**Figure 4.**
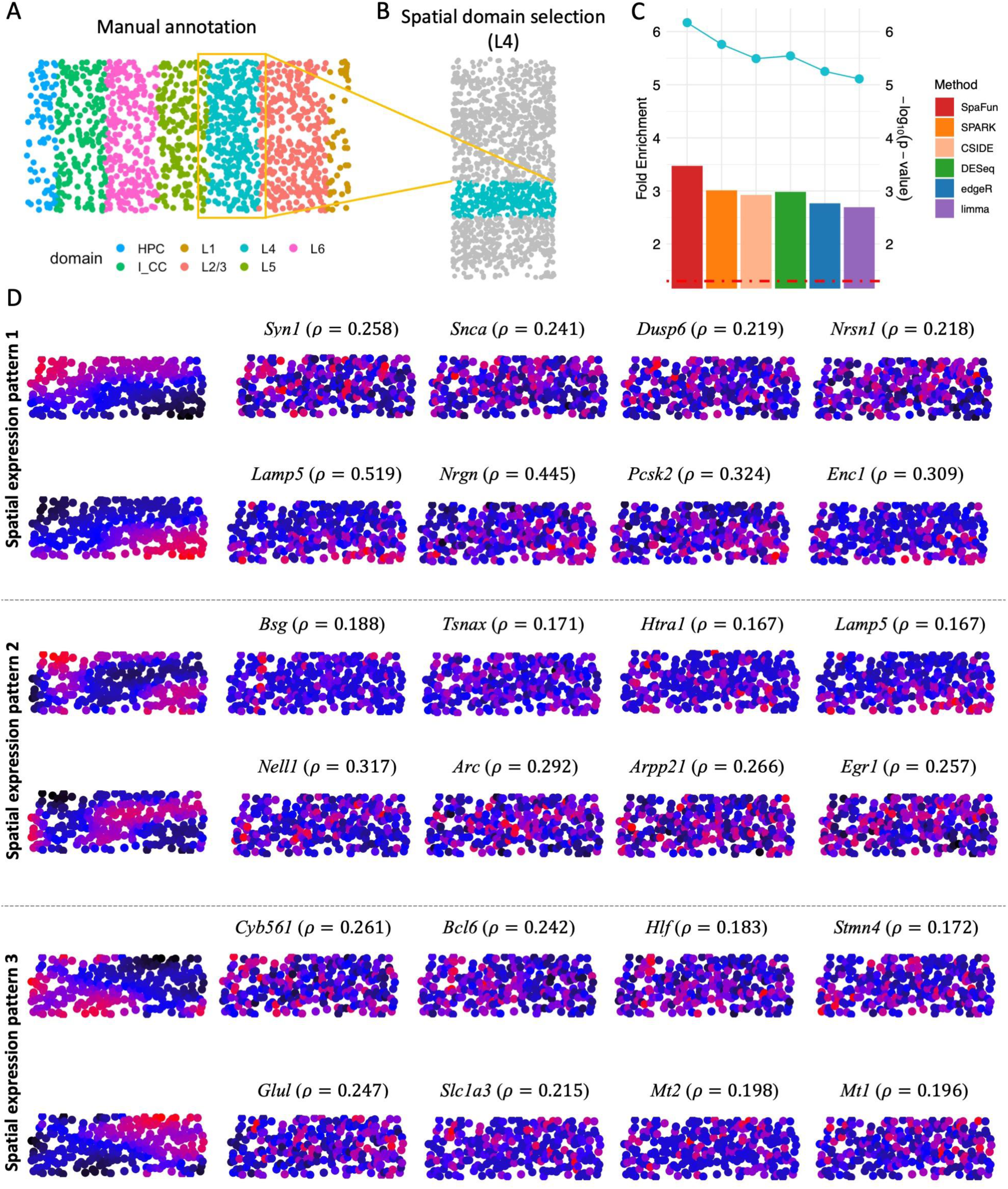
Result of the mouse visual cortex STARmap dataset. (**A**) The pathologist’s annotation of spatial domains. (**B**) Layer L4 selected for analysis. (**C**) Gene set enrichment analysis results for six methods: SpaFun, SPARK, CSIDE, DESeq, edgeR, and limma. Bar plots represent the fold enrichment of each method, with the dotted blue line indicating the −log_10_(p-value) for each method and the dashed red horizontal line marking the threshold where p-value = 0.05. (**D**) The top three principal components (PCs) in layer L4, each shown in two opposite directions, along with the corresponding genes exhibiting similar expression patterns to each PC.

The spatial expression patterns for the top three PCs are shown in Figure 4D. For PC1, the positive direction exhibits higher expression levels in the top-left corner and lower expression levels in the bottom-right corner. Correspondingly, genes Syn1 and Snca share a similar expression pattern with the positive direction of PC1, while Lamp5, Nrgn, Pcsk2, and Enc1 are highly correlated with the negative direction of PC1. PC2 displays a distinct pattern from PC1, with lower expression levels in the middle and higher expression levels in the top-left and bottom-right corners for the positive direction. For PC3, the positive direction shows higher expression levels in the bottom-left corner and lower expression levels in the top-right corner. The genes identified as highly correlated with each PC are presented in Figure 4D.

We also conducted a comparison experiment on six methods (SpaFun, SPARK, CSIDE, DESeq, edgeR, and limma), and the results are presented in Figure 4C. The dotted blue line represents the −log_10_(p-value) for each method, and the dashed horizontal red line is where p-value = 0.05. Four methods (SpaFun, SPARK, edgeR, and limma) obtained a p-value < 0.05, while the other two methods (CSIDE and DESeq) had p-values > 0.05. The accompanying bar plot illustrates the fold enrichment across all the methods. Among them, SpaFun achieved the lowest p-value and the highest fold enrichment, underscoring its effectiveness in identifying biologically meaningful genes in single-cell-level SRT data.

## DISCUSSION

SpaFun offers a groundbreaking approach to detecting domain representative genes (DRGs) in spatially resolved transcriptomics (SRT) datasets, addressing key limitations of existing methods. Traditional SVG detection methods often suffer from high computational complexity, limited statistical power, and an inability to effectively consider spatial heterogeneity and gene co-expression. These limitations are especially pronounced in large-scale datasets where biological and technical variability can obscure meaningful spatial patterns. By leveraging functional principal component analysis (fPCA), SpaFun identifies DRGs with distinct spatial expression patterns specific to annotated regions such as tumor domains, without relying on predefined spatial distribution assumptions.

One of SpaFun’s key advantages lies in its ability to treat gene expression as a function of spatial location. This enables the method to uncover spatially localized expression patterns that are biologically relevant and representative of the underlying tissue architecture. Unlike global SVG detection approaches, SpaFun focuses on domain-specific analysis, capturing localized gene expression dynamics that are often critical for understanding complex biological processes such as tumor microenvironment interactions, tissue heterogeneity, and disease progression.

Applications to three SRT datasets—two from 10X Visium and one from STARmap— demonstrated SpaFun’s robustness and versatility. SpaFun consistently identified DRGs with biologically meaningful spatial patterns in tumor domains, surpassing other methods in statistical power and interpretability. Comparative analyses showed that SpaFun outperformed competing techniques in enrichment analyses and its predicted genes overlapped with known gene sets from the COSMIC database, highlighting its potential to identify disease-relevant genes. For instance, in the human prostate cancer dataset, SpaFun detected distinct spatial patterns in tumor regions. Similar results were observed in the ovarian cancer dataset and the mouse visual cortex dataset, where SpaFun effectively captured both macro-scale spatial heterogeneity and micro-scale co-expression patterns.

Beyond its methodological contributions, SpaFun represents a paradigm shift in how spatial transcriptomics data are analyzed. By integrating spatial heterogeneity and co-expression relationships, it offers a holistic perspective on tissue organization and gene regulation. This capability is particularly relevant for translational research, as understanding localized gene expression patterns can reveal new therapeutic targets and biomarkers for disease diagnosis and treatment. Furthermore, SpaFun’s non-model-based framework ensures scalability to different SRT platforms and experimental designs, making it a versatile tool for diverse applications in spatial biology.

Despite its strengths, SpaFun has limitations that warrant further investigation. For example, the reliance on pre-annotated domains assumes accurate prior knowledge, which may not always be available. Additionally, while SpaFun efficiently handles large datasets, further optimization could improve its computational performance for ultra-large-scale applications. Future work could also explore the integration of histological and molecular profiles to enhance the detection of spatial patterns and extend the applicability of SpaFun to multi-modal data.

In conclusion, SpaFun represents a significant advancement in spatial transcriptomics, offering a robust and scalable approach for detecting spatially variable genes with domain-specific expression patterns. By addressing existing limitations and introducing novel analytical capabilities, SpaFun paves the way for deeper insights into tissue biology, disease mechanisms, and potential therapeutic interventions. Its application across diverse datasets highlights its utility and sets a strong foundation for future developments in spatially resolved data analysis.

## DATA AVAILABILITY

An open-source implementation of the SpaFun algorithm in R/C++ is available at https://github.com/yg2485/SpaFun. The authors analyzed three publicly available SRT datasets. Raw count matrices, images, and spatial data for two SRT datasets from 10x Visium are accessible on the 10x Genomics website at https://support.10xgenomics.com/spatial-gene-expression/datasets. Mouse visual cortex STARmap data can be downloaded from https://www.starmapresources.com/data.

## References

1. Wang X, Allen WE, Wright MA et al. Three-dimensional intact-tissue sequencing of single-cell transcriptional states, Science 2018;361.

2. Chen KH, Boettiger AN, Moffitt JR, et al. RNA imaging. Spatially resolved, highly multiplexed RNA profiling in single cells, Science 2015;348:.

3. Lubeck E, Coskun AF, Zhiyentayev T et al. Single-cell in situ RNA profiling by sequential hybridization. Nat Methods. 2014, 360–361.

4. Femino AM, Fay FS, Fogarty K et al. Visualization of single RNA transcripts in situ, Science 1998;280:585–590.

5. Raj A, Van Den Bogaard P, Rifkin SA et al. Imaging individual mRNA molecules using multiple singly labeled probes, Nature Methods 2008;5:877–879.

6. Lubeck E, Coskun AF, Zhiyentayev T et al. Single-cell in situ RNA profiling by sequential hybridization, Nature Methods 2014;11:360–361.

7. Chen KH, Boettiger AN, Moffitt JR et al. Spatially resolved, highly multiplexed RNA profiling in single cells, Science 2015;348:.

8. Ståhl PL, Salmén F, Vickovic S et al. Visualization and analysis of gene expression in tissue sections by spatial transcriptomics, Science 2016;353:78–82.

9. Rao N, Clark S, Habern O. Bridging genomics and tissue pathology: 10x genomics explores new frontiers with the visium spatial gene expression solution, Genetic Engineering & Biotechnology News 2020;40:50–51.

10. Rodriques SG, Stickels RR, Goeva A et al. Slide-seq: A scalable technology for measuring genome-wide expression at high spatial resolution, Science 2019;363:1463–1467.

11. Yip SH, Sham PC, Wang J. Evaluation of tools for highly variable gene discovery from single-cell RNA-seq data, Briefings in Bioinformatics 2019;20:1583–1589.

12. Leek JT, Storey JD. Capturing heterogeneity in gene expression studies by surrogate variable analysis, PLoS genetics 2007;3:e161.

13. Zhao R, Lu J, Zhou W et al. A systematic evaluation of highly variable gene selection methods for single-cell RNA-sequencing, BioRxiv 2024:2024.

14. Svensson V, Teichmann SA, Stegle O. SpatialDE: identification of spatially variable genes, Nature Methods 2018;15:343–346.

15. Sun S, Zhu J, Zhou X. Statistical analysis of spatial expression patterns for spatially resolved transcriptomic studies, Nature Methods 2020;17:193–200.

16. BinTayyash N, Georgaka S, John S et al. Non-parametric modelling of temporal and spatial counts data from RNA-seq experiments, Bioinformatics 2021;37:3788–3795.

17. Li Q, Zhang M, Xie Y et al. Bayesian modeling of spatial molecular profiling data via Gaussian process, Bioinformatics 2021;37:4129–4136.

18. Vandenbon A, Diez D. A clustering-independent method for finding differentially expressed genes in single-cell transcriptome data, Nature Communications 2020;11:4318.

19. Miller BF, Bambah-Mukku D, Dulac C et al. Characterizing spatial gene expression heterogeneity in spatially resolved single-cell transcriptomic data with nonuniform cellular densities, Genome research 2021;31:1843–1855.

20. Dries R, Zhu Q, Dong R et al. Giotto: a toolbox for integrative analysis and visualization of spatial expression data, Genome Biology 2021;22:1–31.

21. Hu J, Li X, Coleman K et al. SpaGCN: Integrating gene expression, spatial location and histology to identify spatial domains and spatially variable genes by graph convolutional network, Nature Methods 2021;18:1342–1351.

22. Wong RK, Zhang X. Nonparametric operator-regularized covariance function estimation for functional data, Computational statistics & data analysis 2019;131:131–144.

23. Wang J, Wong RK, Zhang X. Low-rank covariance function estimation for multidimensional functional data, Journal of the American statistical Association 2022;117:809–822.

24. Fukumizu K, Gretton A, Lanckriet G et al. Kernel choice and classifiability for RKHS embeddings of probability distributions, Advances in neural information processing systems 2009;22.

