## Supplementary Information for "SpaFun: Discovering Domain-specific Spatial Expression Patterns and New Disease-Relevant Genes using Functional Principal Component Analysis"

### for

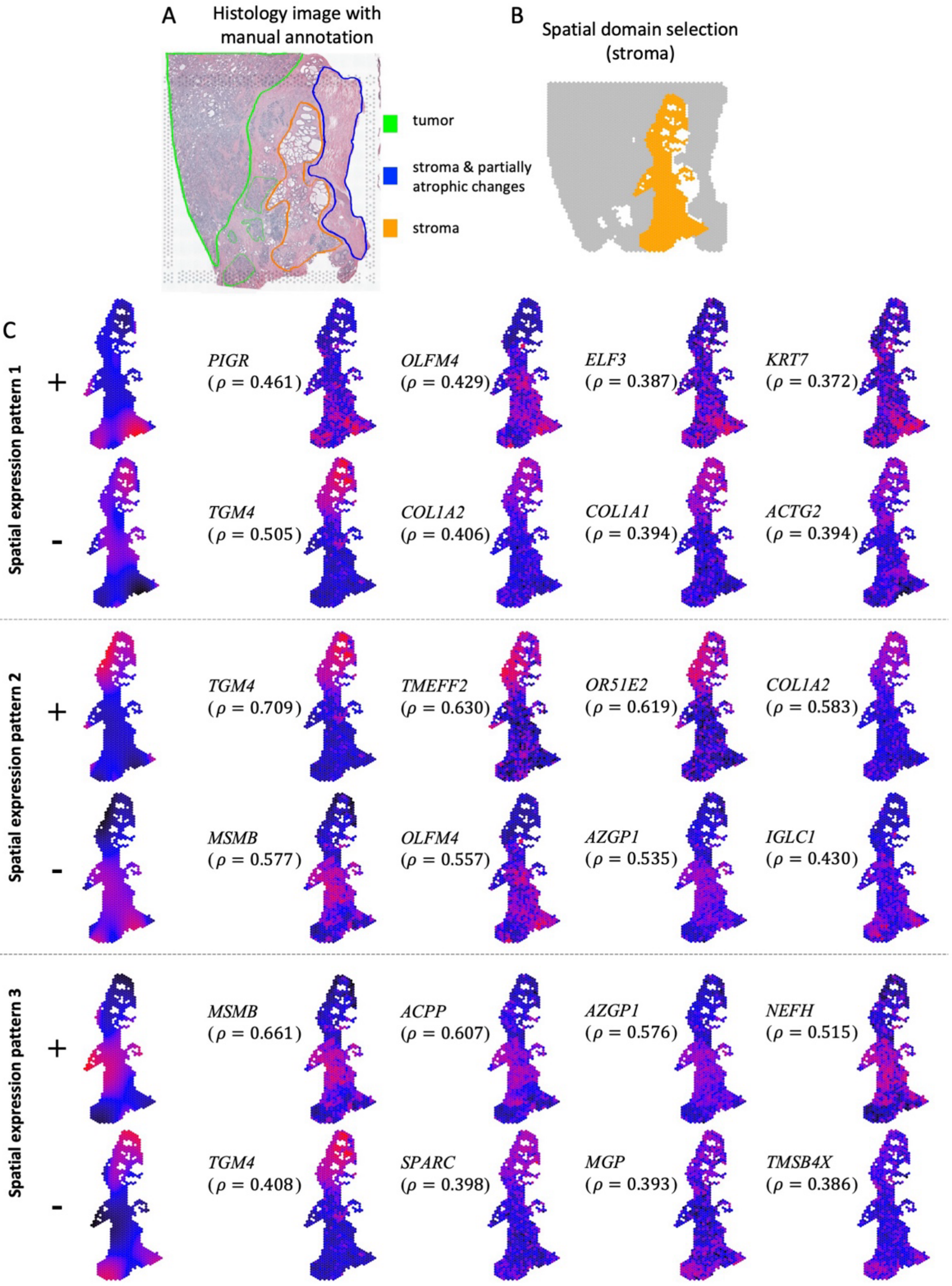


Figure S1. Additional results of the human prostate cancer dataset. (**A**) Histology image with pathologist-annotated spatial domains. (**B**) Stroma domain selected for analysis. (**C**) The top three principal components (PCs) in the tumor domain, each shown with two opposite directions, along with the corresponding genes exhibiting similar expression patterns to each PC.

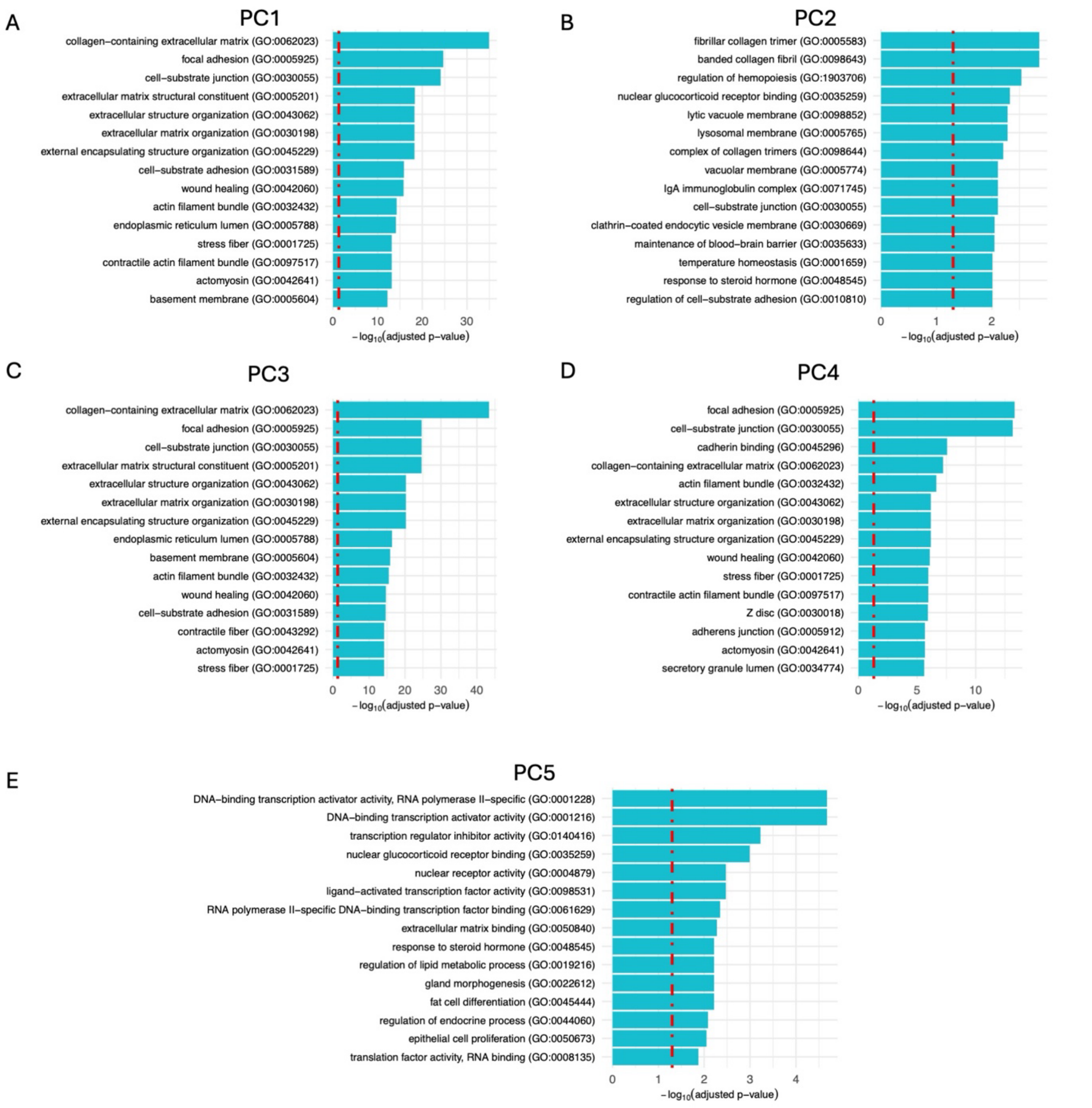


Figure S2. Gene Ontology (GO) enrichment analysis on the human prostate cancer dataset. The analysis is conducted on the selected DRGs for the first five PCs. The first 15 GO terms with smallest p-values are displayed, with the dashed red line representing the threshold where p-value = 0.05.

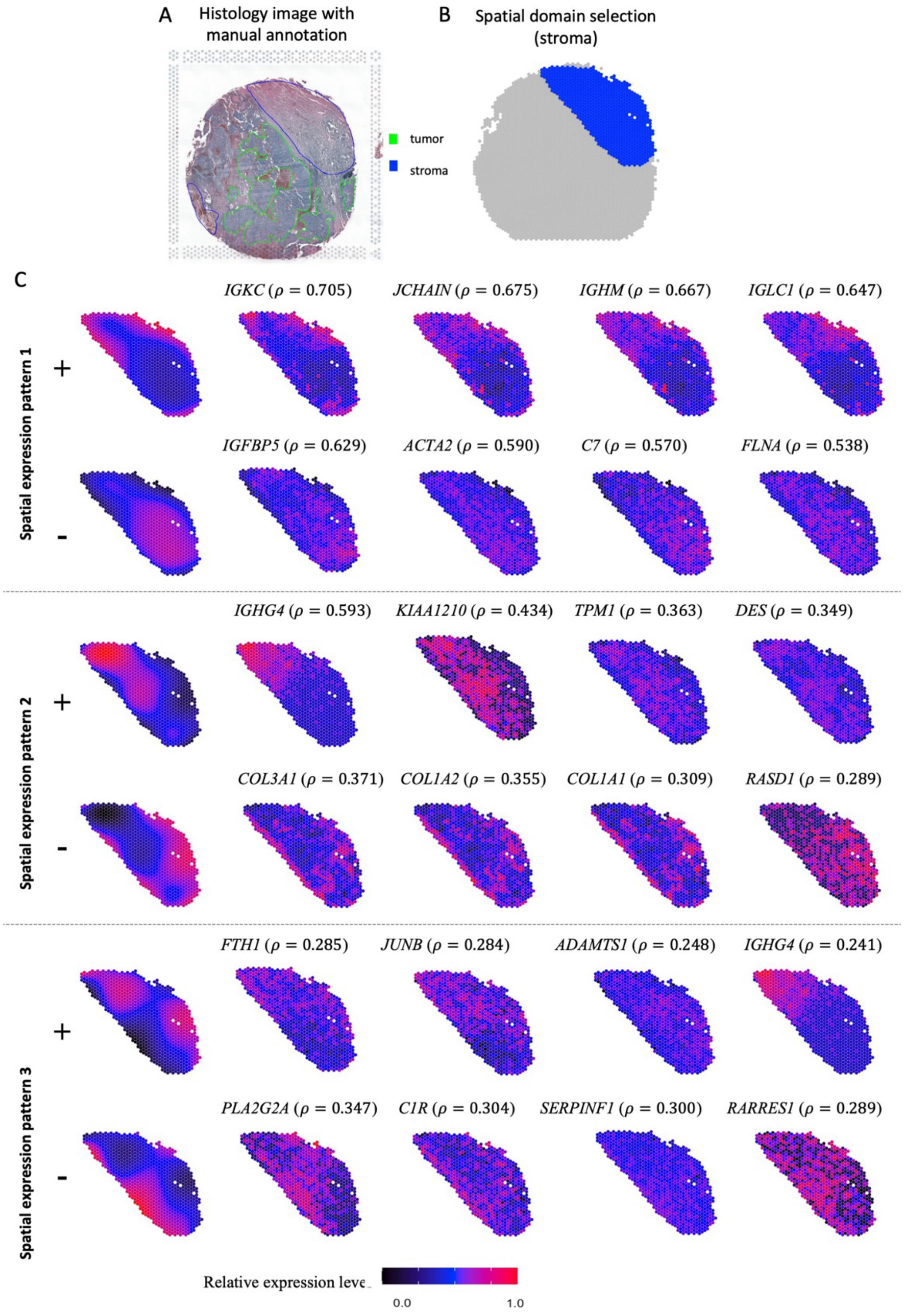


Figure S3. Additional result of the human ovarian cancer dataset. (**A**) Histology image with pathologist-annotated spatial domains. (**B**) Stroma domain selected for analysis. (**C**) The top three principal components (PCs) in the tumor domain, each shown with two opposite directions, along with the corresponding genes exhibiting similar expression patterns to each PC.

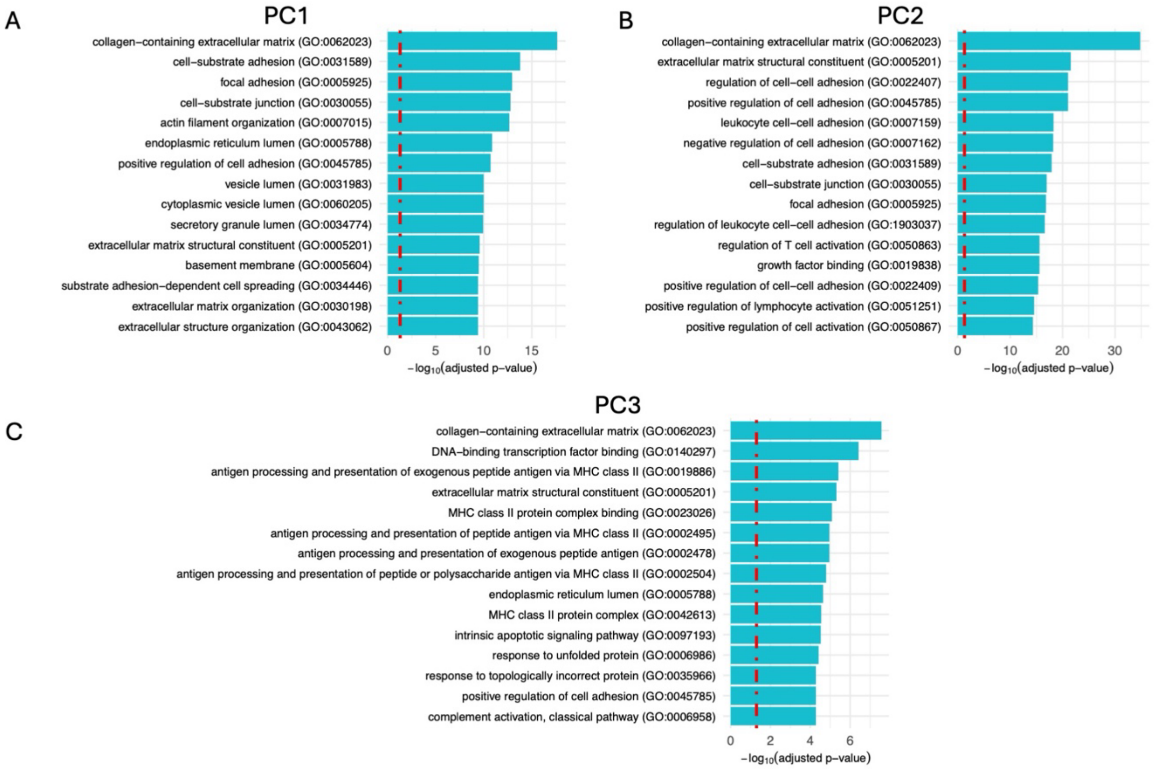


Figure S4. Gene Ontology (GO) enrichment analysis on the human ovarian cancer dataset. The analysis is conducted on the selected DRGs for the first three PCs. The first 15 GO terms with smallest p-values are displayed, with the dashed red line representing the threshold where p-value = 0.05.

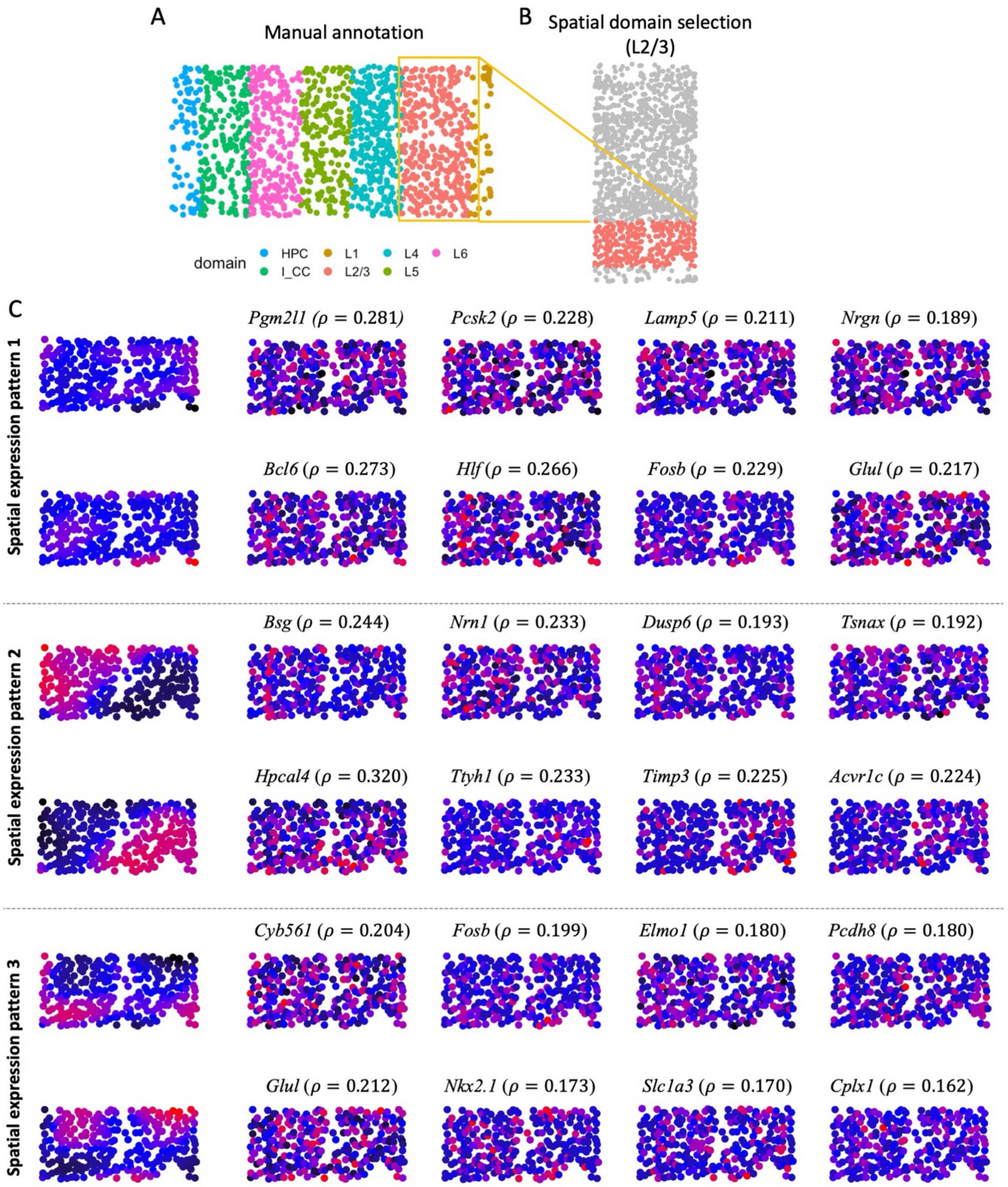


Figure S5. Additional result of the mouse visual cortex STARmap dataset. (**A**) Pathologist's annotation of spatial domains. (**B**) Layer L2/3 selected for analysis. (**C**) The top three principal components (PCs) in layer L2/3, each shown with two opposite directions, along with the corresponding genes exhibiting similar expression patterns to each PC.
